# Natural variants in SARS-CoV-2 S protein pinpoint structural and functional hotspots: implications for prophylaxis and therapeutic strategies

**DOI:** 10.1101/2021.01.04.425340

**Authors:** Suman Pokhrel, Benjamin R. Kraemer, Scott Burkholz, Daria Mochly-Rosen

## Abstract

In December 2019, a novel coronavirus, termed severe acute respiratory syndrome coronavirus 2 (SARS-CoV-2), was identified as the cause of pneumonia with severe respiratory distress and outbreaks in Wuhan, China. The rapid and global spread of SARS-CoV-2 resulted in the coronavirus 2019 (COVID-19) pandemic. Earlier during the pandemic, there were limited genetic viral variations. As millions of people became infected, multiple single amino acid substitutions emerged. Many of these substitutions have no consequences. However, some of the new variants show a greater infection rate, more severe disease, and reduced sensitivity to current prophylaxes and treatments. Of particular importance in SARS-CoV-2 transmission are mutations that occur in the Spike (S) protein, the protein on the viral outer envelope that binds to the human angiotensin-converting enzyme receptor (hACE2). Here, we conducted a comprehensive analysis of 441,168 individual virus sequences isolated from humans throughout the world. From the individual sequences, we identified 3,540 unique amino acid substitutions in the S protein. Analysis of these different variants in the S protein pinpointed important functional and structural sites in the protein. This information may guide the development of effective vaccines and therapeutics to help arrest the spread of the COVID-19 pandemic.

## Introduction

To curb the COVID-19 pandemic, most efforts have focused on preventing entry of the virus by inhibiting the interaction of severe acute respiratory syndrome coronavirus 2 (SARS-CoV-2) with its human receptor, angiotensin-converting enzyme 2 (hACE2)^1^. Interaction of SARS-CoV-2 with hACE2 occurs via the S protein on the viral envelope. Proteases cleave the S protein into S1 and S2 subunits^2,3,4^ to enable viral binding to hACE2^5^ and viral entry by membrane fusion^6^. The S protein is a homotrimer and the S1 subunit of each of the monomers of the S protein contains the receptor-binding domain (RBD; Fig. 1a, b) in either the ‘open’ (active) or ‘closed’ (inactive) conformations^7,8,9^ (Supplemental Fig. 1a).

**Figure 1.**
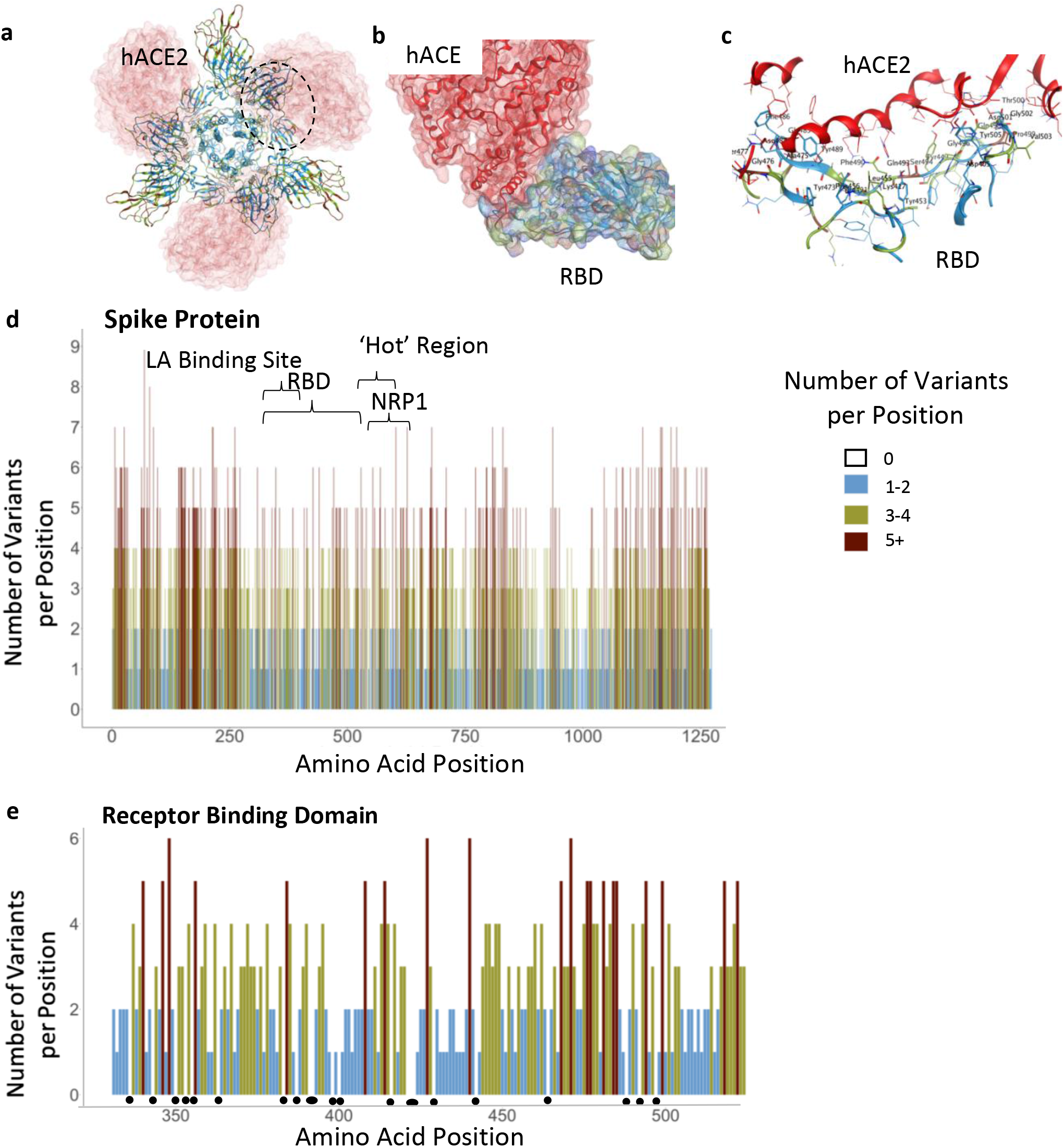
Functional regions in S protein and the RBD-hACE2 interaction site. **a)** S protein homotrimer with ribbons colored according to legend, bound to hACE2 (red). Black dotted outline shown in **b. b)** RBD-hACE2 interface. **c**) RBD-hACE2 interface highlighting residues in RBD within 4.5Å from hACE2. **d)** The number of variants per position across the entire sequence of S protein, highlighting specific functional regions. **e)** The number of variants per position across RBD. Black dots indicate invariable positions.

Four main types of prophylaxis or therapeutic strategies, focusing on the S protein, have been employed: 1). Preventing proteolysis of the S protein^10^; 2). Competing with S1 binding to hACE2, using S1 or hACE2 protein fragments or peptides^1,11,12^; 3). Generating monoclonal or polyclonal antibodies against SARS-CoV-2 S protein or RBD, to be used as passive vaccines^13^; and 4) Active vaccines that generate an immune response, usually to the S1 subunit^14,15,16,17^.

Besides the RBD, the S protein of the coronaviruses, including SARS CoV-2, has several other regions that are predicted to be relatively conserved due to their critical role for S protein functions. These regions include the trimer interface of S^7,9^, furin proteolysis sites^5,6^, glycosylation sites^18,19^, neuropilin-binding sites^20,21,22^ and linoleic acid (LA)-binding site^9,23^. These regions may be important for maintaining structural integrity, entry, and transmission of the virus and therefore are likely to serve as potential targets for development of prophylaxes and therapeutics.

Although SARS-CoV-2 undergoes mutations at a lower frequency than other viruses like influenza and HIV^24^, the emergence of several common variants of SARS-CoV-2 in human populations may generate resistance to current prophylaxis and therapeutics. Some of these mutations result in gain of fitness for the virus due to mutations in the S protein^25,26,27,28^. Early in the pandemic, in February 2020, a single missense mutation resulting in a change from aspartate to glycine in position 614 (D614G) emerged in Europe and became the dominant variant of the virus. The D614G variant has spread throughout the world and increased the transmissibility of SARS-CoV-2 by conferring higher viral loads in young hosts without an apparent increase in the severity of the disease^29^. With the emergence of new variants, such as B.1.1.7 (also known as the UK variant) and B. 1.351 (also known as the South African variant) that have greater transmissibility and escape antibody detection^25,26,27,28^ (Table 1), it is imperative to map other substitutions in the S protein sequence. Such substitutions may contribute to future variants that lead to increased transmissibility or to variants that evade prophylaxis or therapeutics. Particularly, amino acid substitutions in the RBD, including those that interact directly with hACE2^25,26,27,28^ (Fig. 1c) may have an impact. Here, we aimed to identify regions on the S protein that are relatively invariant to guide prophylaxis and therapeutic development more efficiently.

## Results

### SARS-CoV-2 Spike Protein

The SARS-CoV-2 S protein is 1273 amino acids long; it contains a signal peptide (amino acids 1-13), the S1 subunit (14-685 residues) that mediates receptor binding, and the S2 subunit (686-1273 residues) that mediates membrane fusion^30^. To identify areas in the S protein that are the least divergent as the virus evolves in humans, we obtained viral sequences from GISAID (Supplemental Table 1) that as of March 1, 2021, included 633,137 individual virus sequences isolated from humans throughout the world. As compared with the index WIV04 (MN996528.1, also known as the Wuhan variant or index virus) sequence of February 2020^31^, the 1,273 amino acid S protein^8^ had 3,540 variants. This number of variants only includes filtered sequences (441,168) that are complete and do not contain an abnormal number of mutations (see Methods). As there are 3,540 variants, on average, each position in the 1273 amino acid protein sequence has approximately three variants (Fig. 1d). However, some regions harbor 9 variants in a single amino acid position whereas others have no variants (Fig. 1d; Supplemental Table 3). Regions in S protein with 2 or fewer variants/position (marked in light blue, Fig. 1d, e) are more prevalent in the structurally critical trimer interface (46% of the amino acids; Fig. 1d, Supplementary Fig. 1b, c, see Supplementary Table 4), and in the RBD (56%, Fig. 1e, Supplementary Fig. 1b, c). There are a total of 123 positions that are entirely invariable (Supplementary Table 3).

### Receptor Binding Domain

Much of the prophylaxis and therapeutic efforts are focused on the RBD (amino acids 331-524). Among the 3,540 variant sequences, we found only 22 invariant amino acids in the RBD (Fig. 1e, marked by dots under the position; Supplemental Table 3). Of those amino acid substitutions in the RBD, only 3% are predicted by PROVEAN software^32^ to be structurally or functionally damaging (Supplemental Table 2). Using PROVEAN, we also examined the predicted impact of the amino acid substitutions in the common more infective variants (B.1, B.1.1.7, B.1.351, B.1.427/429, B.1.256 and P.1; Table1) on the RBD structure and function and found that these variants are predicted to have a neutral effect, suggesting these variants are not decreasing the fitness of the virus.

### Furin Proteolysis Sites

We next examined other regions in the S protein for which functions have been assigned. Furin proteolysis at the S1-S2 boundary (681-685) and in S2 (811-815) exposes the RBD to enable hACE2 binding, and the S2 domain to initiate membrane fusion^5^. Recent studies show that these cleavage sites are not necessarily specific for furin-mediated proteolysis and that S may be processed by multiple proteases to open the RBD into the active conformation^2,3,4,33^. Consistent with these observations, both the furin proteolysis consensus sites and the arginine that are critical for proteolysis are not conserved in the S protein (Fig. 2a), in agreement with a prior analysis of furin site 1^34^.

**Figure 2.**
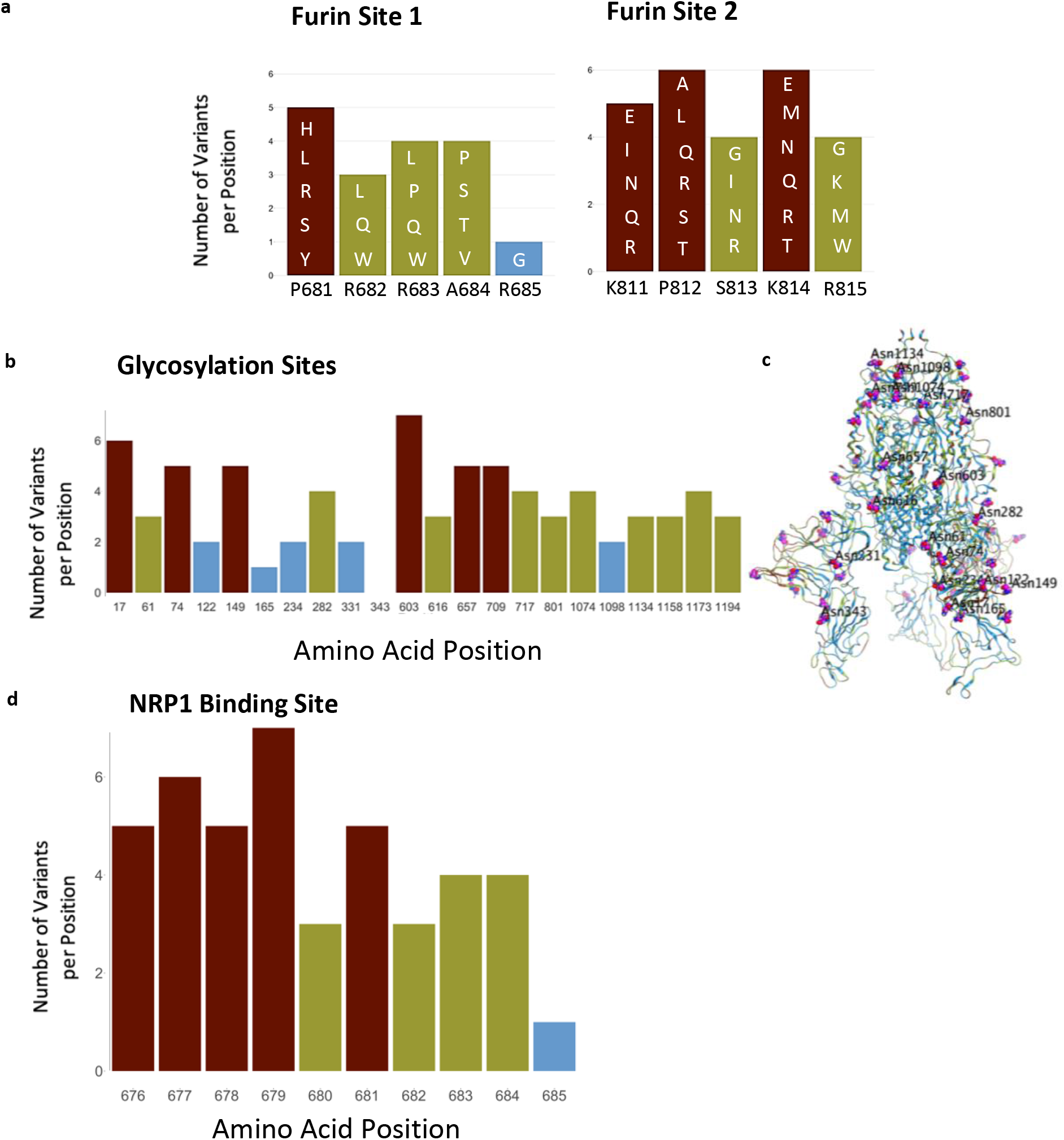
Furin cleavage sites, glycosylation sites, and NRP-1 interaction site. **a)** Furin cleavage sites in the S protein. Amino acid indicated below each bar indicate the sequence in the WIV04 index isolate. Variants in other SARS-CoV-2 isolates are indicated within the bars of the graphs using one letter abbreviations for the amino acids. **b)** The 22 glycosylation sites in the S protein; indicated are the number of variants per position. **c)** Glycosylation asparagine sites (numbered) are highlighted in pink in the S protein. **D)** The number of variants in the proposed NRP1-binding site.

### Glycosylation Sites

The S protein also has 66 glycosylation sites in each trimer, which facilitate protein folding and may lead to host immune system evasion^19^, as 40% of the S protein’s surface is shielded by glycans^18^. Surprisingly, with one exception, none of these glycosylation sites were invariable, suggesting that not all the glycosylation sites are essential for the S protein’s functions (Fig. 2b,c). The only asparagine serving as an invariable glycosylation site is N343 in the RBD domain, located more than 25Å away from hACE2-binding site, and therefore unlikely to mediate receptor binding.

### Neuropilin-1 Interaction Site

Neuropilin-1 (NRP-1) is a transmembrane receptor that regulates angiogenesis^35^ and immune response^36^ and is expressed in many cell types^36^ such as the endothelium^37^, immune cells^38^, and neurons^39^. Interaction between NRP-1 and S protein was proposed to regulate SARS-CoV-2 transmission^20,21,22^. Proteolysis of S1 index variant by furin was found to expose a C terminal motif, RXXR (where R is arginine and X is any amino acid), known to be the binding motif in NRP-1^20,22^. For example, a monoclonal antibody against the RXXR-binding site on NRP-1 SARS-CoV-2 reduced infectivity in culture^22^. Nevertheless, we found that the NRP-1 interaction-site in S1 is not conserved (Fig. 2d). Although the variants are predicted to have a neutral effect on the S1 protein structure (using PROVEAN analysis, Supplemental Table 2), 90% of the positions in the NRP1-interaction site have more than 2 variants (or an average of 4.3 variants/position; Fig. 2d).

### Linoleic Acid-Binding Site

A fatty acid-binding pocket has been identified in the inactive conformation of S protein^9^ (Fig. 3a, b). The amino acids that make this pocket are conserved in other coronaviruses^9^ and are unchanged (less than 2 variants) in 75% of the position (Fig. 3a, b). Furthermore, among the 20 amino acids that line this pocket, 71% of the identified variants are predicted to have a neutral effect using PROVEAN (Supplemental Table 2). Analysis of the LA-bonding site identifies a potential pharmacophore that may fit small molecules (Fig. 3c), perhaps by mimicking ω-3 fatty acids^23^.

**Figure 3.**
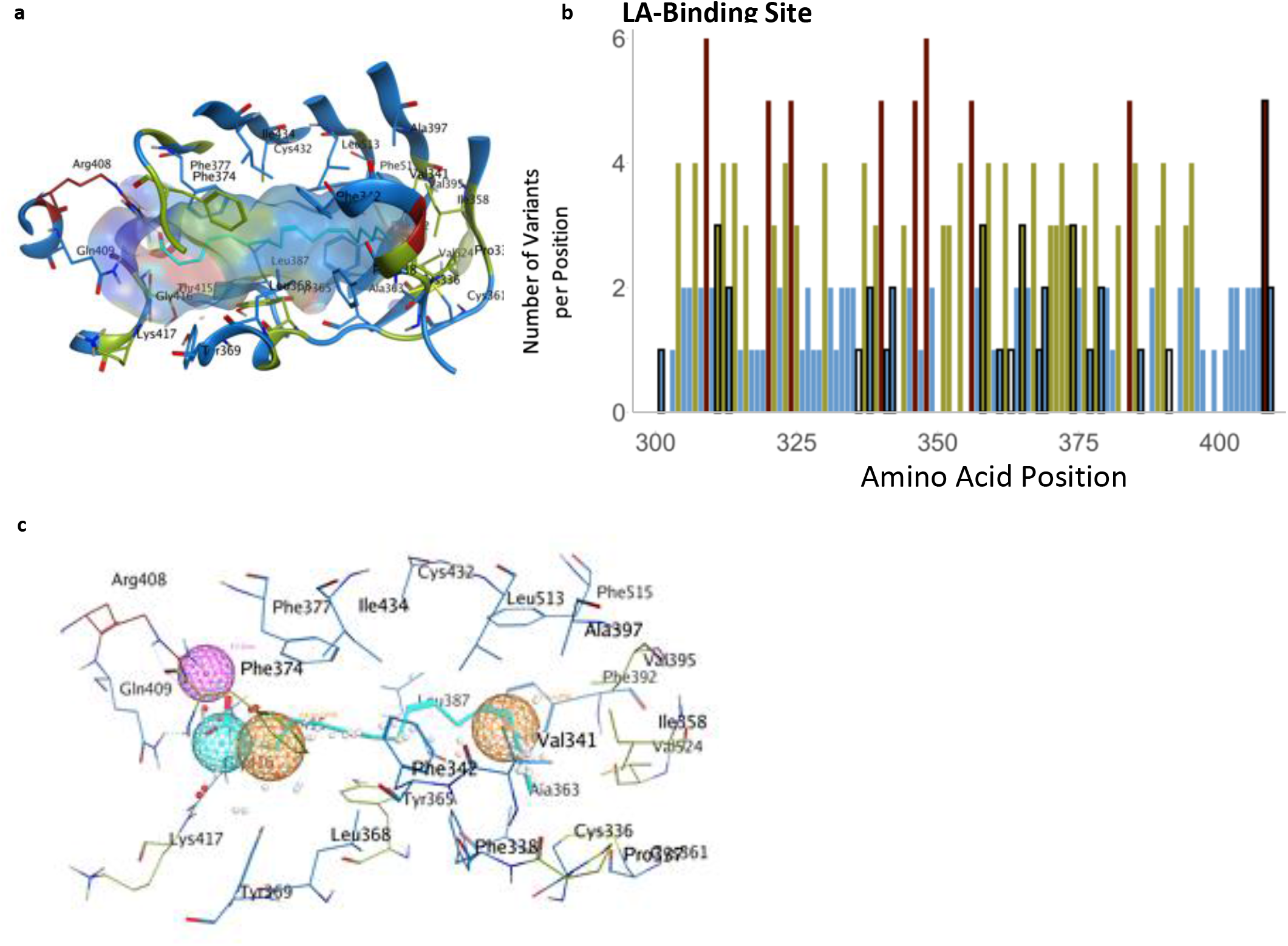
The LA-binding site in the S protein. **a)** Hydrophobic pocket forms the LA-binding site. **b)** The number of variants per position across the LA-binding site; black outlines indicate the positions that form the LA-pocket. **c)** Pharmacophore of the LA-binding pocket.

### Relatively Invariable Regions with Unidentified Function

We also identified another less variable region between residues 541-612 (Fig. 4); 62% of the amino acid positions in this region have 2 or fewer variants and 12 positions are entirely invariable (‘Hot Region’; Fig. 1a and Fig. 4a, b). This less variable region is relatively hydrophobic, yet a substantial number of residues remain exposed in the open and closed conformations (Fig. 4c). Six residues, V551, T553, C590, V595, V608, Y612, in this relatively invariable region form a part of the largest hydrophobic patch in the protein measuring 370 Å^2^ (Fig. 4d, e). Five of these residues (excluding T553) along with other residues that make this hydrophobic patch tolerate very few mutations and almost all the mutations that are tolerated change to hydrophobic amino acids (Fig. 4d). We examined this region using Site Finder in MOE^40^ and found that there is a binding site with a positive score for the propensity of ligand binding^41^, which encompasses several residues from this region (i.e. Cys590, Ser591, Phe592, Gly593) (Supplemental Fig. 1e). This hydrophobic region is also 81% identical between SARS-CoV and SARS-CoV-2, but less than 15% identical when comparing the SARS-CoV-2 sequence with that of MERS-CoV (Fig. 4f).

**Figure 4.**
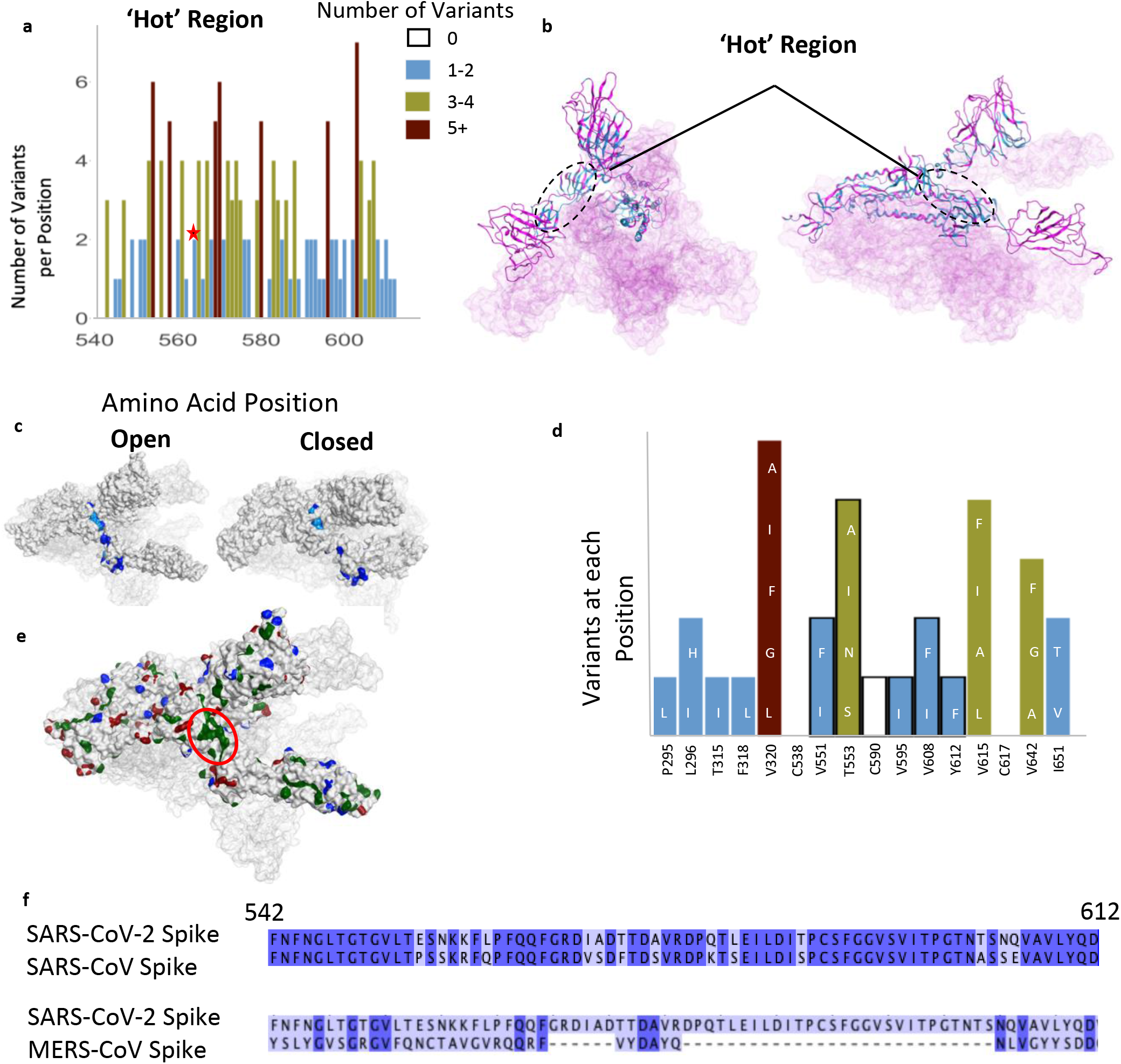
A relatively invariant (‘hot’) region in the S protein with no known function, identified by analyzing 441,168 individual virus sequences. **a)** The number of variants per position across the less-variable, ‘hot’ region with un-assigned function. The red star identifies the proposed ‘latch’, Q564 residue. **b)** The hot region identified in the 3-D structure of S (open conformation). **c)** Invariant ‘hot’ region in S protein with un-assigned function depicted in both the open (left) and closed (right) conformations. Dark blue denotes invariant amino acids and light blue denotes positions with 1-2 observed variants. This region becomes exposed after S protein gets activated by proteases. **d)** Number of variants in hydrophobic patch with unassigned function. Positions outlined in black are part of the ‘hotspot’. **e)** Some residues in the hotspot (shown in **d**) are part of the largest hydrophobic patch (green, red ellipsoid) of S protein. Positive patches are highlighted in red. Negative patches are highlighted in blue. **f**) Sequence identity between SARS-CoV-2 & SARS-CoV (81% identical), and SARS-CoV-2 & MERS-CoV (15%) in the ‘hotspot’. Dark blue denotes identical amino acid residues. Numbering corresponds to SARS-CoV-2.

## Discussion

While SARS-CoV-2 has a lower mutation rate than other viruses due to proof-reading mechanisms^24^, aspects such as a relatively high R_0_ of 1.9 to 2.6^42^, comparatively long asymptomatic incubation and infection periods, and zoonotic origins, leads to high variability in mutations in specific regions compared to the original reference sequence. This has been illustrated with the divergence of 6 major lineages in the past few months (Table 1). Our analysis of the frequency of variants throughout the S protein of SARS-CoV-2 identified regions of high and low divergence, which may aid in developing effective prophylactic and therapeutic treatments. In this analysis of mutations in the S protein, we did not consider the frequency of a particular mutation or in how many countries the mutation was found. Such analysis, as was done for D614G^43^, may further aid in determining the potential improved viral fitness acquired by a particular mutation.

**Table 1.**
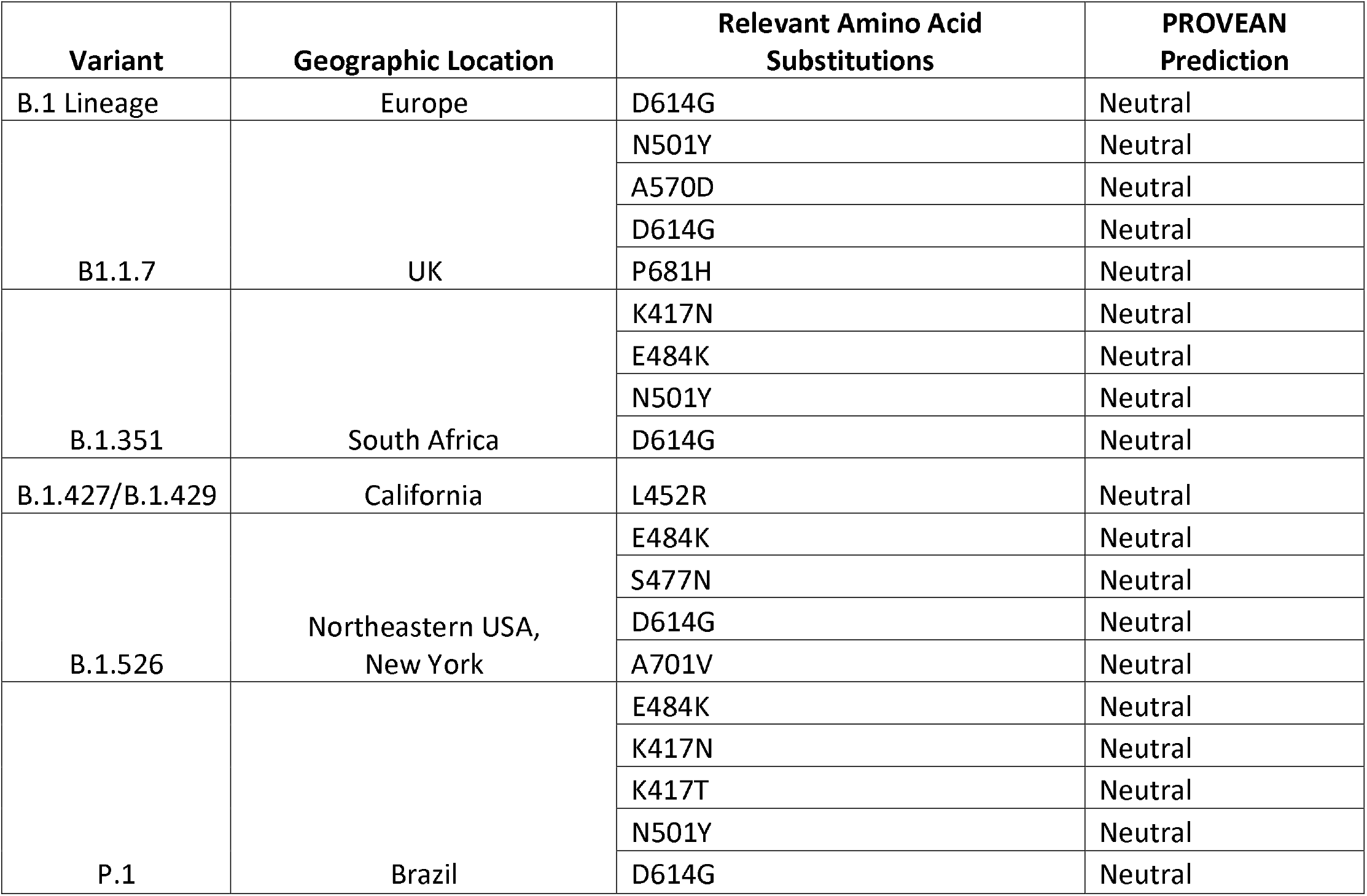
Common variants of concern.

Protein glycosylation is essential for viral infection.^44^ In SARS-CoV-2 S protein, there are 22 known N-glycosylation sites per monomer (Fig. 2b, c), but only one, asparagine 343, appears to be conserved. Furthermore, we found 156 positions in S that mutate to an asparagine residue in the existing 3,540 variants that we analyzed, and many of them are exposed on the S protein (Supplemental Fig. 1d). We propose that some of these new asparagine residues may create new glycosylation sites on the S protein that can contribute to immune evasion. Such an impact on the immune evasion by changes in the positions of glycosylation sites of viral envelope proteins have been described for influenza viruses; e.g., H3N2 has numerous new N-linked glycans on the viral hemagglutinin that enabled the virus to escape antibody neutralization and evade the host’s immune system^45^. The formation of new glycosylation positions may also affect viral susceptibility to existing antibodies and to the immune response of infected individuals. A cryo-electron microscopy study has already suggested that coronaviruses mask important immunogenic sites on their surface by glycosylation^46^. Furthermore, recent work suggests that changes in glycosylation sites may affect its recognition by other potential human proteins and receptors, inducing the toll-like receptors, calcitonin-like receptors, and heat shock protein GRP78, thus leading to a more severe inflammation that characterizes a more severe form of COVID-19^47^.

Additional sites on the S protein have been suggested to be critical for viral infectivity, including the trimer interface, the furin proteolysis sites and the NRP-1 binding site. However, our analysis suggests that not all these sites will be effective targets for prophylaxis and therapeutics. Specifically, the trimer interface is less accessible and therefore unlikely to be druggable. Another issue relates to the furin sites. As the viral S protein activation appears to require furin proteolysis^2,3,4^, protease-specific inhibitors are tested as a means to protect from infection^48^. However, our analysis suggests that this may not be an effective strategy, given the high variability of furin cleavage sites. This suggestion is consistent with previous data showing that other proteinases expressed throughout the body may work synergistically to activate the S protein^2,33^. Therefore, drugs that focus on inhibiting any single protease may not be effective preventative treatment against all SARS-CoV-2 variants. Similarly, the NRP1-binding site that is generated by proteolysis and the exposure of a C-terminal RXXR motif^20,22^ may not be a good target for treatment against all SARS-CoV-2 variants, unless such a motif is created by other proteases.

Are there additional sites on the S1 protein that can be explored to identify new treatments of COVID-19 or prevention of infections by SARS-CoV-2? There might be a benefit in focusing on the LA-binding site that help stabilize the S in the inactive closed conformer. Small molecules that mimic LA and bind into the LA pocket may stabilize the S protein in the closed/inactive conformation, thus reducing infectivity (Fig. 3). Therefore, exploring the LA pharmacophore (Fig. 3c) with small molecules that can hold the S-protein in closed conformation, thus preventing the presentation of RBD to hACE2, could be of great interest as this may reduce viral infectivity. Our data also suggest that it may be beneficial to develop passive and active vaccines that target the RBD, instead of the entire glycosylated S protein; the RBD is less variable relative to the whole S1 protein (compare Fig. 1e to 1d). However, similar to some of the common viral isolates, such as the South African, B. 1.351, new amino acid substitutions in the RBD may evade such therapeutics; e.g., loss of immunoreactivity to monoclonal antibodies^25^.

Finally, our study suggests that drugs and antibodies targeting region 541-612, a relatively conserved and exposed region on the protein’s surface that we identified (Fig. 4), warrant further study. Determining how druggable the pocket encompassing this region is (residues Cys590, Ser591, Phe592, Gly593), provided its solvent exposure, and whether modulating S protein by engaging this site will have a biological consequence is a challenge (Supplemental Fig. 1e). Very recently, Q564 within this region (star in Fig. 4a) has been proposed to act as a ‘latch’, stabilizing the closed/inactive conformation of the S protein^49^. The high degree of conservation of hydrophobicity in this region potentially indicates its role in membrane fusion and/or maintaining structural integrity. The sequence similarity between SARS-CoV-2 and SARS-CoV (Fig. 4f) further supports the importance of this region, especially as both viruses have a similar route of infection. Determining the role of this invariable region warrant a further study, as it may be another Achilles heel to target for anti-SARS-CoV-2 treatment.

## Materials & Methods

### Database of S protein amino acid variants, the world regions from where the virus was obtained, and whether the sequence is predicted to be deleterious

A FASTA formatted file containing 633,137 spike protein sequences was retrieved on 03/01 from the GISAID database. This file had previously been preprocessed by the database with the individual alignment of genomes to the WIV04 (MN996528.1^31^) reference sequence, using mafft [https://doi.org/10.1093/molbev/mst010], *via* the command “mafft --thread 1 --quiet input.fasta > output.fasta” with subsequent translation into protein from the spike protein-coding region at 21563 to 25384.

For the analysis in this paper, only sequences sampled from humans were retrieved with the spike protein sequences realigned *via* mafft [https://doi.org/10.1093/molbev/mst010] against the WIV04 (MN996528.1, http://dx.doi.org/10.1038/s41586-020-2012-7) reference utilizing parameters ideal for a large number of highly similar protein sequences as well as using the option to maintain position numbering against the reference.

“grep -i “|Human|” input.fasta -A1 > output.fasta”

“mafft --6merpair --thread −1 --keeplength --addfragments input.fasta reference.fasta > output.fasta”

A python script (supplementary material) was generated to filter sequences based on set quality thresholds that included 1) 0 ambiguous protein positions; 2) 0 deletions or gaps outside of common deletions including position 69, 70 and 144/145; 3) only full length pre-alignment of 1273 but down to 1270 in the event of the specified deletions; and 4) a maximum of less than 1% (13) amino acid substitutions from reference. These resulting 441,168 sequences (supplementary table), were chosen by the strict quality thresholds to remove low quality and potentially error prone sequences based on those that were incomplete, contain uncommon deletions, insertions, and have an unusually high number of mutations.

### Calculating Number of Variants

The raw data for variants in the S protein (see Extended Material Table) was read into R studio (v. 1.3.1093) and analyzed using the Tidyverse package. The number of unique variants was calculated for each position, excluding insertions. Graphs were created for specific regions and each position was color-coded according to the number of variants present in that position (i.e., 0 – no color, 1-2 is blue, 3-4 is yellow, > 5 is red). See sample code below:

~~~
Calculating variants:
df%>%
 group_by(Position, .drop=FALSE)%>%
 tally()
Graphing Example:
ggplot(df)+ #graph of RBD, works for diff colors
 geom_col(data = subset(df, Position > 330 & Position < 525), aes(x=Position, y=(n), fill=as.factor(n)))+
 ggtitle(“RBD”) +
 scale_fill_manual(values = pal, name = “Number”) +
 labs(y = “Number of Mutations”)+
 theme(panel.background = element_blank(), text=element_text(size=20))
~~~

For the functional regions, the proportion of positions with 2 or fewer observed variants was calculated. See formula below:

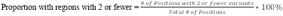

### Calculating Predicted Effect of Variants in PROVEAN

The amino acid sequence of S protein from the reference EPI_ISL_402124 (WIV04; Wuhan) sequence was uploaded to PROVEAN (http://provean.jcvi.org/index.php). Every variant observed in S was also uploaded to compare against the reference sequence. Each variant was either predicted to be ‘deleterious’ or ‘neutral’. The PROVEAN predictions were also read into R studio (v. 1.3.1093) and analyzed with the Tidyverse package for every region analyzed. The proportion of variants predicted to be neutral and deleterious were calculated for the functional regions analyzed in S. See Supplemental Table 1. Sample code below:

~~~
Calculating PROVEAN ratios:
table(df$ProveanPrediction)%>%
 prop.table()%>%
 round(4)
~~~

### Protein Structures

Molecular Operating Environment (MOE) software^40^ was used to prepare the figures using PDB ID: 7A98^4^ for Figures 1b-d, 2d; Supplemental Fig. 1a (left), d (left), Supplemental Fig. 2a, and PDB ID: 6ZB5^6^ was used to prepare Fig. 2a, Supplemental Fig. 1a (right), d (right).

### Sequence Alignment

The Spike protein sequences from SARS-CoV-2, SARS-CoV, and MERS-CoV were uploaded to Jalview^50^. The Mafft alignment was then performed to align each amino acid sequence.

### Pharmacophore Generation

PDB ID: 6ZB5^6^ was opened and prepared using the QuickPrep functionality at the default settings in MOE as mentioned previously. Dummy atoms were created at the LA-binding site formed by chains 6ZB5.A and 6ZB5.C. AutoPH4 tool^51, 52^ was used to generate the pharmacophore at the dummy atom site in the Apo generation mode.

## Acknowledgments

Supported in part by the 2020 COVID-19 Response: Drug and Vaccine Prototyping Grant from the Innovative Medicines Accelerator, Stanford University to D. M.-R. We gratefully thank the many investigators throughout the world that provided the SARS-CoV-2 sequences to this public database.

## Author Contributions

S.P., B.R.K., S.B. provided data analysis, visualization, and draft writing. D. M.-R. conceived the project, supervised the analysis and writing.

---

**Supplementary figure 1.**
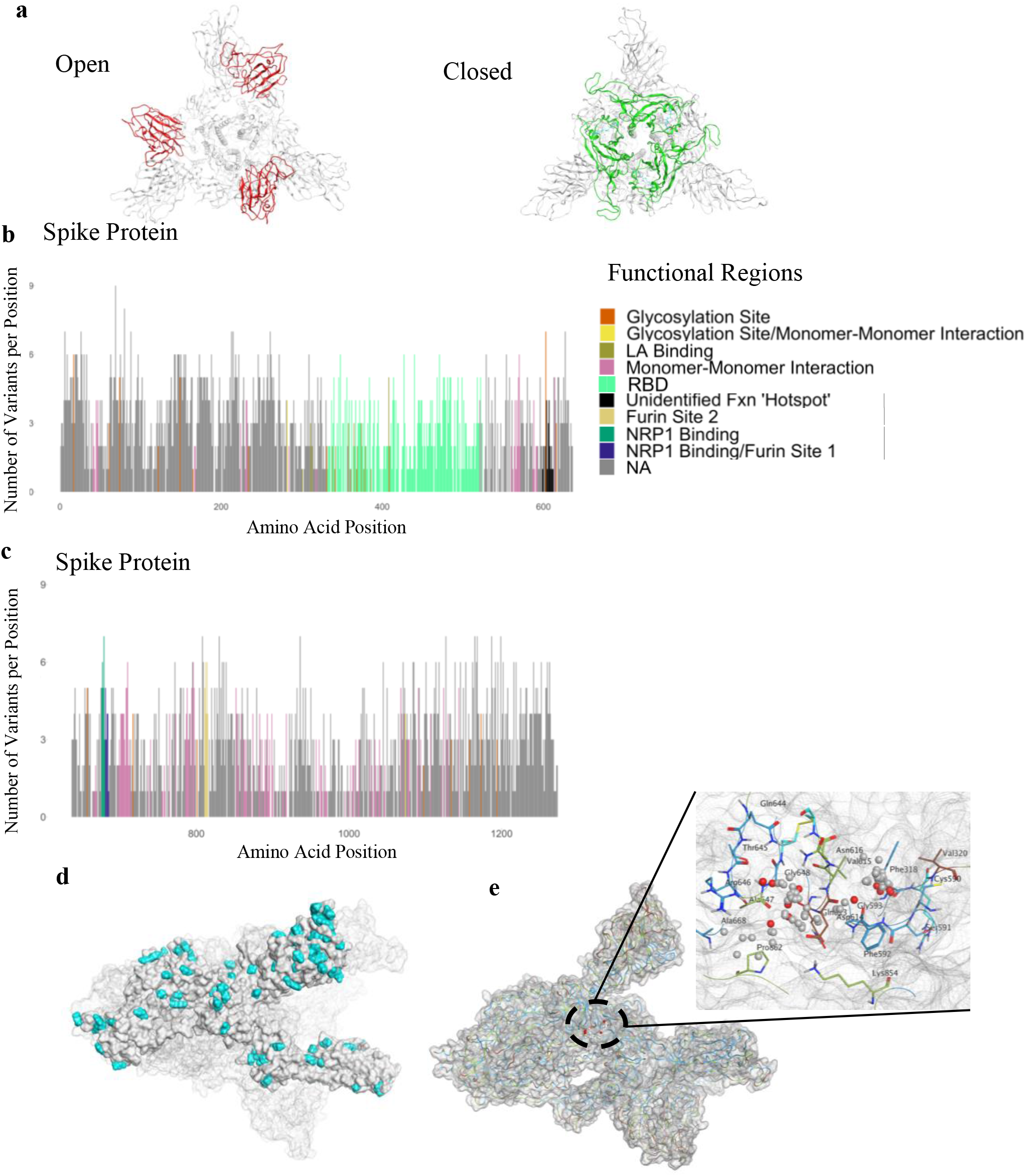
Variant and invariant regions in SARS-CoV-2 protein. **a)** SARS-CoV-2 Spike (S) protein trimer showing the receptor-binding domain (RBD) in the open conformation (red; PDB ID: 7A98) and the closed conformation (green; PDB ID: 6ZB5). Viewed from the top. **b & c**) Regions in the sequence (1-636: **b**; 637-1273:**c**) of the S protein are color-coded according to function. **d)** New potential glycosylation sites (cyan) on S protein in circulating variants. **e)** Potential ligand binding site in the hotspot (aa 541-612).

